# Transcriptional Differences of Peanut (Arachis hypogaea L.) Seeds in the Fresh-harvest, After-ripened and Just-germinated States: insights into regulatory networks of seed dormancy-release and germination

**DOI:** 10.1101/690248

**Authors:** Pingli Xu, Guiying Tang, Weipei Cui, Guangxia Chen, Chang-Le Ma, Jieqiong Zhu, Pengxiang Li, Lei Shan, Zhanji Liu, Shubo Wan

## Abstract

Seed dormancy and germination are the two important traits related to plant survival and reproduction, and crop yield. To understand their regulation mechanism, it is crucial to clarify which genes or which pathways participate in the regulation of these processes. However, little information is available during the procedure of seed dormancy and germination in peanut. In this study, the seeds of the variety Luhua No.14 with non-deep dormancy were selected and its transcriptional changes at three developmental stages: the fresh-harvest (FS), the after-ripened (DS) and the just-germinated seeds (GS), were investigated by comparative transcriptomics analysis. The results showed that genes with increased transcription in DS vs FS comparison were overrepresented for oxidative phosphorylation, glycolysis pathway and tricarboxylic acid cycle (TCA), suggesting that after a period of drying storage, the intermediates stored in dry seeds were rapidly mobilized by glycolysis, TCA cycle, glyoxylate cycle, etc.; the electron transport chain accompanying with respiration has been reactivated to provide ATP for mobilization of other reserves and seed germination. In GS vs DS pairwise, dozens of the up-regulated genes were related to plant hormone biosynthesis and signal transduction, including the majority of components in auxin signal pathway, and brassinosteroid biosynthesis and signal transduction, and some GA and ABA signal transduction genes. During seeds germination, the expression of some *EXPANSIN* and *XYLOGLUCAN ENDOTRANSGLYCOSYLASE* was also significantly enhanced. To investigate the effect of different hormone during the procedure of seed germination, the contents and the differential distribution of ABA, GA, BR and IAA in cotyledon, hypocotyl and radicle, and plumule of three seed sections at different developmental stages were also detected. Combining with previous data in other species, a model of regulatory network related to peanut seed germination was developed. This model will helpful to gain further understanding of the mechanisms controlling seed dormancy and germination.

## Introduction

Seed dormancy and germination are the two important traits in the plant life cycle, which involve in the survival of a species and the offspring proliferation. Different plant species have various classes of dormancy to regulate the timing of seed germination, help seedlings emerge under favorable conditions. Primary dormancy of seeds is acquired during the seed maturation phase, and reaches a high level in freshly harvested seeds. During subsequent dry period of seeds (after-ripening), primary dormancy slowly reduces. When the dormancy level gradually decreases, seeds can rapidly loose dormancy and proceed to germination during imbibition at favorable conditions [1]. A recent research in *Arabidopsis* suggested that seed after-ripening is a specific developmental pathway that is independent of germination potential and doesn’t rely on ABA regulation [2]. The dormancy alleviation in dry seeds is associated with ROS production and the carbonylation of specific embryo proteins [3–5]. Concomitantly, the metabolic switches between different developmental periods of seeds are also relevant to the distinct expression profiles of genes involved in several metabolism pathways [6].

Strictly defined, germination is the initial emergence of the radicle from the seed coat. In some species, like as *Arabidopsis*, whose embryo is enclosed by the endosperm and the surrounding testa, seed germination consists of two visible steps: first, the testa rupture due to expansion of the endosperm and embryo, followed by the radicle protruding through the endosperm. However, in leguminous plants, seed is endospermless, and testa splitting marks the completing germination [7–9]. Phytohormones play the important roles in the induction and the maintenance of seed dormancy, as well as the release of dormancy and the following germination. Abscisic acid (ABA) and gibberellins (GA) negatively and positively regulate seed germination. In different development states of seed, the ratio between ABA and GA in embryo is changeable. The dormant seeds maintain a high ABA/GA ratio, and dormancy maintenance also depends on high ABA/GA ratios, while dormancy release involves a net shift to increased GA biosynthesis and ABA degradation resulting in low ABA/GA ratios, and seed germination associated with the increasing of GA content and sensitivity [7,10]. The basic role of auxin is to promote cell elongation. Increasing of GA content leads to the obvious change in auxin content and transport during seed germination. A peak of free IAA appears prior to the initiation of radicle elongation [11]. Brassinosteroid (BR), as another antagonist of ABA, and GA play the parallel roles to promote cell elongation and germination. Photodormancy is released by the GA/light signal transduction pathway, while the subsequent endosperm rupture is activated by the BR and the GA/light pathways with distinct mechanisms [11–13].

In recent years, many studies have focused on gene expression analyses related to seed dormancy and germination, and have revealed some genes that regulate seed dormancy and germination, especially genes involved in phytohormone signaling such as ABA, GA, BR and IAA pathways [9,14–22]. Many transcriptome analyses involved in seed dormancy and germination in different plants describe a global view of gene expression changes among different developmental stages, or at different region of seeds, or in dormant and non-dormant seeds [15,23–25]. Bassel et al. (2011) indicated that the characters of seed dormancy and germination are much conservative in evolution of flowering plants. The genome-wide transcriptional analyses of dormant and after-ripened *Arabidopsis* seeds over four time points and two seed compartments found that the gene sets strongly enhanced at the initiation of imbibition are overrepresented for GO classes including key cellular metabolic processes like translation, amino acid, organic acid, nucleotide and carbohydrate metabolism, and the down-regulated sets includes response to stress and other environmental cues [15]. During germination of soybean seeds, GA, ethylene and BR pathways are transcriptionally active, while ABA signalling is down-regulated in the embryonic axes [26].

Cultivated peanut (*Arachis hypogaea* L.) is a distinctive oilseeds crop, which flowers on the ground and fructifies under the ground. After peanut maturity, harvest delayed may cause *in situ* germination of seeds when they meet constant rainy days, always leading to a depression in yield and a reduction in seed quality. The species *A. hypogaea* L. has been divided into two subspecies: *A. hypogaea subsp. hypogaea* and *A. hypogaea subsp. fastigiata*. In the subspecies *A. hypogaea subsp. hypogaea var. hypogaea* (Virginia and Runner market types) and *var. hursita*, varieties have longer growth cycle and seeds have longer dormancy stage. While in subspecies *fastigiata* involving *var. fastigiata* (Valencia market class) and *var. vulgaris* (Spanish market type), varieties are early-maturing but generally lack fresh seed dormancy [27]. However, even the *subsp. hypogaea* with dormancy, their dormant status is easy to be broken during storing for a short time at room temperature. In our present research, in order to explore the regulatory mechanism of dormancy release and germination of peanut seeds, the seeds of the variety Luhua No.14 (LH14) belonging to *subsp. hypogaea* with non-deep dormancy of seeds were selected and its transcriptional changes at three developmental stages: the fresh-harvest, the after-ripened and the just-germinated seeds, were investigated by comparative transcriptomics analysis.

## Materials and Methods

### Plant Material and Growth Conditions

Peanut variety LH14 bred by Shandong Peanut Research Institute, and planted by our group in the field located at Yinmaquan Farm for subsequent assay and analysis. The seeds were harvested from the field and divided into two parts, one portion of them harvested freshly was kept in paper bags under −80°C or under ambient temperature and humidity, and another part designation as after-ripened seeds was dried in sunshine for over two weeks and kept in paper bags under room temperature. For the assay of germination rate, 5 accessions [2 from *subsp. fastigiata*: chico (CHI) and Silihong (SLH); 3 from *subsp. hypogaea*: LH14, Fenghua No.1 (FH1) and Linguimake (LGMK)] were selected. Thirty six seeds from each accession were sowed in three petri dishes over four layers of absorbent gauze wetted with demineralized water and incubated in 15°C incubator with darkness. The status of imbibition was determined at 24 h intervals based on the changes of seed swelling, and seeds were considered germinated when the radicles broke through seed coat.

### RNA Extraction, Library Construction and Sequencing

The whole seeds from fresh harvest (FS), after ripened (DS), and germinated exactly (GS) were collected for RNA extraction. Two biological repeats were set up. Total RNA of the samples was isolated using the improved CTAB method [28], and was treated with DNase I (RNase-free) according to the TaKaRa’s protocol. Their quantity and purity was measured using Qubit^®^ RNA Assay Kit in Qubit^®^ 2.0 Flurometer (Life Technologies, CA, USA) and the NanoPhotometer^®^ spectrophotometer (IMPLEN, CA, USA), and the integrity was examined with the RNA Nano 6000 Assay Kit of the Bioanalyzer 2100 system (Agilent Technologies, CA, USA).

A total amount of 3μg RNA each sample was used to construct cDNA library. The libraries were generated using NEBNext^®^ Ultra™ RNA Library Prep Kit for Illumina^®^ (NEB, USA) following manufacturer’s recommendations. Sequencing for 6 libraries was completed by Beijing Novogene Bioinformatics Technology Co., Ltd. on a HiSeq 2000 platform (Illumina, San Diego, CA, USA), and 100 bp paired-end reads were generated.

### Data Analysis for RNA-Seq

The raw data of fastq format were cleaned by removing adapter sequences, reads containing poly-N, and low-quality reads (Q≤20). The clean reads were aligned to the reference genome (*Arachis ipaensis*, https://www.peanutbase.org/home) using TopHat v2.0.12. All sequence data (Bioproject_accession:PRJNA545858) were submitted to the BioProject database of the National Center for Biotechnology Information (NCBI). The expression levels were calculated with Cufflinks and normalized by the Fragments Per Kilobase of transcript per Millions mapped reads (FPKM) method [29].

The differentially expressed genes (DEGs) between the two samples were detected by DESeq R package (ver. 1. 18. 0) and *P* value were adjusted using the Benjamini and Hochberg method for control of the false discovery rate [30]. Genes with an adjusted *P* < 0.05 were used as the thresholds to determine significant differences in gene expression. Annotation of gene function was performed by comparisons with non-redundant nucleotide and protein sequences (NCBI), and protein sequence database Swiss-Prot. The GO (Gene Ontology) and KEGG (Kyoto Encyclopedia of Genes and Genomes) enrichment analyses were performed to identify which DEGs were significantly enriched in GO terms or metabolic pathways by the GOseq R package and the KOBAS software. GO terms with corrected *P* value less than 0.05 were considered significantly enriched by differential expressed genes. The GO annotations were functionally classified by WEGO software for gene function distributions. The genes with at least twice level of expression in specific developmental stage than that in other stage were defined as preferential DEGs in KEGG pathways referenced by soybean.

### Real-time Quantitative RT-PCR (qRT-PCR) Analysis

DEGs preferentially expressed in specific metabolism pathway and hormone signal pathways were selected for validation using real-time qRT-PCR. The candidate genes and their amplification primers were listed in Table 1. The qRT-PCR was performed by the instruction of SYBR Premix Ex Taq (Tli RNaseH Plus; Takara Biotechnology, Dalian, China). The reaction condition was: predenaturing for 5 min at 94°C, and then 40 cycles of 15 s at 94°C and 30 s at 60°C. The relative expression levels of the target genes were analyzed with *AhACTIN7* as an internal control and calculated using the 2^−ΔΔCt^ method [31].

**Table 1.**
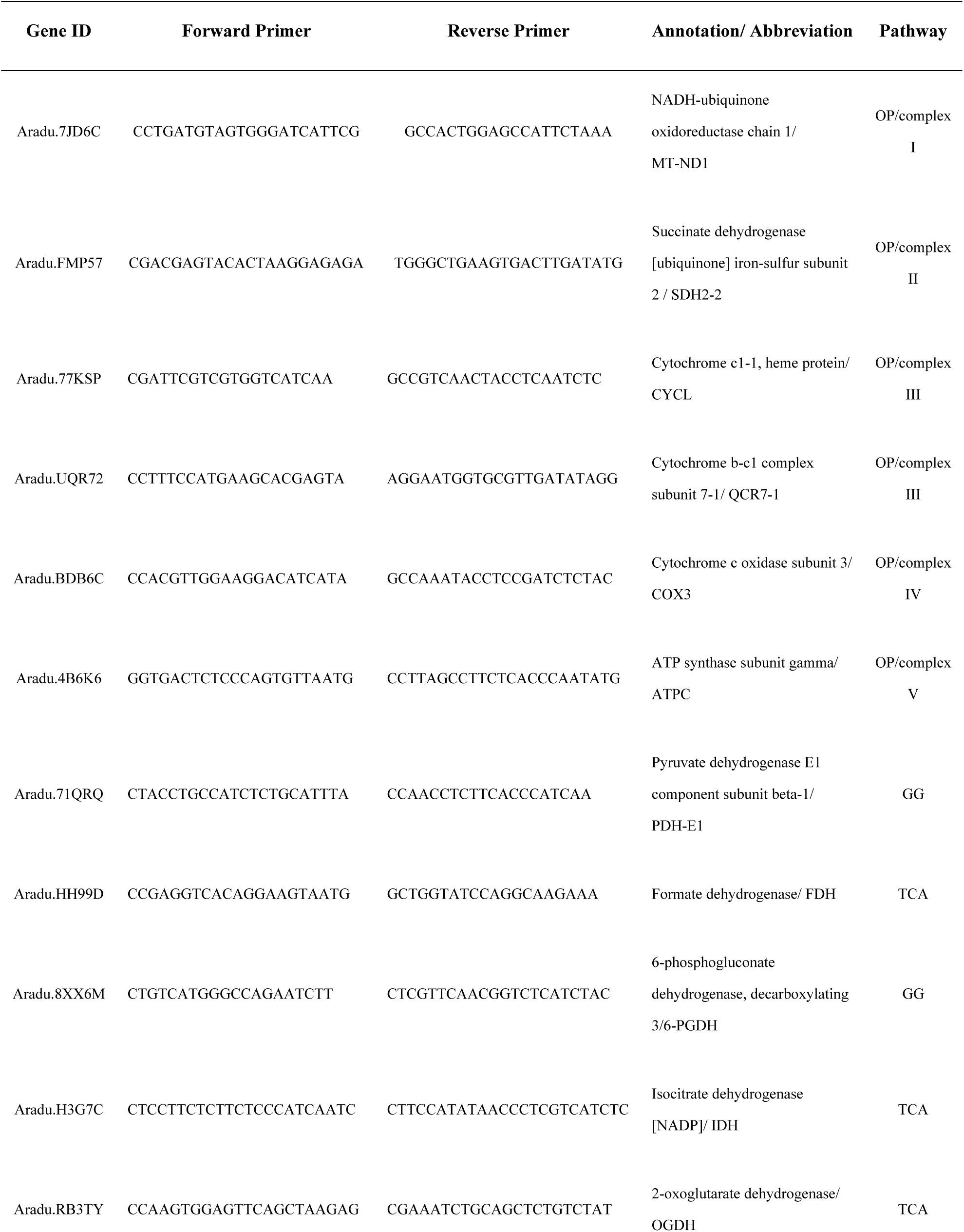
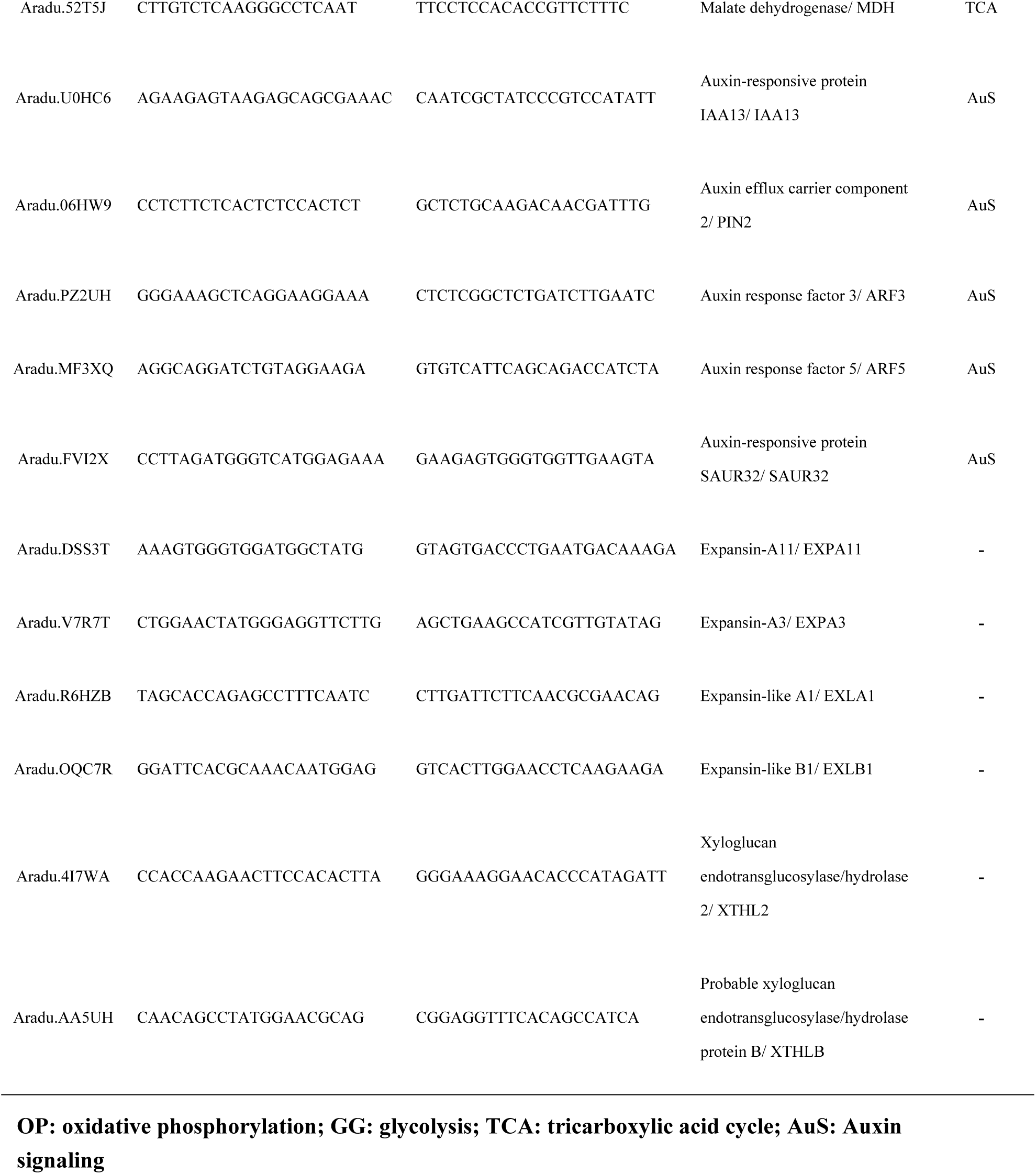
The primers using for verification of gene expression level by qRT-PCR

### Hormones Extraction and Quantification

1g of samples were ground with a mortar and pestle in liquid nitrogen, and extracted in cold 80% (v/v) methanol with butylated hydroxytoluene (1 mmol·L^−1^) overnight at 4°C. The extracts were collected after centrifugation at 10000g for 20 min at 4°C, and purified through a C_18_ Sep-Park Cartridge (Waters Crop., Millford, MA) and dried in N_2_. The hormone fractions were dissolved in phosphate buffer saline (PBS, 0.01 mol·L^−1^, pH7.4) with 0.1% (v/v) Tween 20 and 0.1% (w/v) gelatin for determining the levels of hormones by ELISA.

The peanut seeds were dissected into three parts: cotyledon (CO), hypocotyl and radicle (HR), and the remainder plumule (PL). The contents of ABA, GA_3_, IAA and BR in these three parts were detected by ELISA according to the method reported by Yang et al. [32]. The monoclonal antibodies against ABA, GA_3_, IAA and BR produced by the Phytohormones Research Insititute, China Agricultural University, were used as the first antibody, and IgG horseradish peroxidase was the secondary antibody. The content of each hormone was calculated by known amounts of standard hormone added in the extracts according to description by Weiler et al. [33].

## Results

### Germination Assay

Five accessions including LH14, FH1, LGMK, CHI and SLH, which divided into two classes of non-deep dormant and non-dormant, were selected for the detection of germination rate. To explore the influence of dried storage on seed germination, the germination assay of the fresh-harvest seeds and dry seeds stored for more than two weeks were performed. It was found that the fresh-harvested seeds of LH14, FH1 and LGMK with non-deep dormancy began to germinate after sowing in water for 4d and their germination rates only reached 75%, 77.8%, and 91.7% during imbibiting for more than 10d; and then after a period of dried storage, their majorities of dried seeds started to germinate after uptaking water for 2d and their germination rates reached 100%, 97.2%, and 97.2% after germinated for 4~6d; while in non-dormant peanut seeds from CHI and SLH, there were no obvious difference in the beginning time of germination between the fresh-harvested seeds and the dry seeds, their seeds began to germinate at the second day, but the uniformity of seeds germination in the dry seeds was much higher than that in the fresh-harvested seeds (Fig 1). It was suggested that non-deep dormant peanut seeds released the dormancy through the process of after-ripening by dried storage, whereas non-dormant peanut seeds were short of the typical after-ripening procedure. Thus, peanut variety LH14 with non-deep dormancy was selected using for the following experiments.

**Fig 1.**
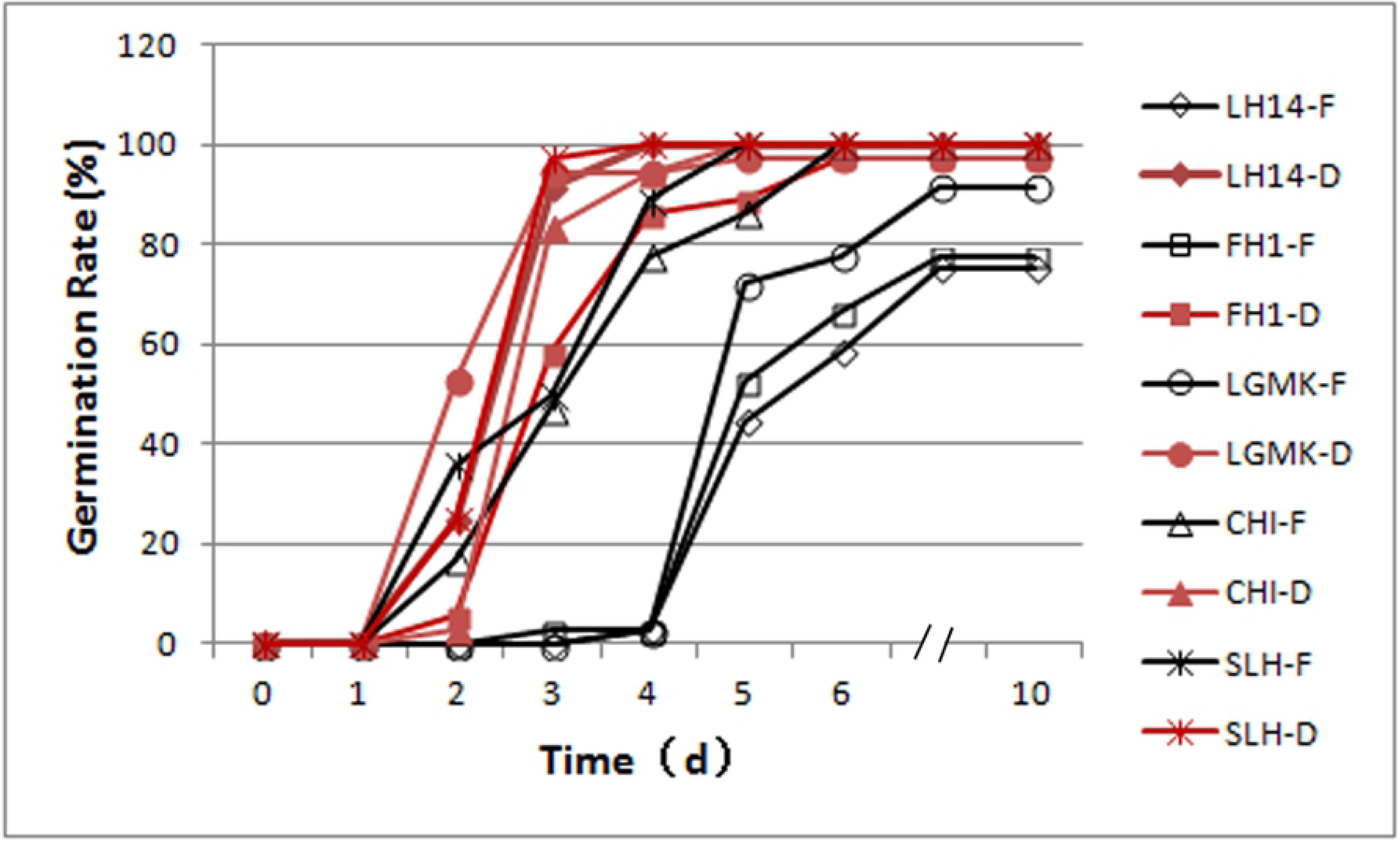
The germination rate of peanut seeds of fresh-harvest (F) and after drying (D) from different varieties LH14: Luhua No. 14, FH1: Fenghua No. 1, LGMK: Lingguimake, CHI: Chico, SLH: Silihong

### Transcriptome Sequencing and Assessment

In order to investigate the changes of transcript profile in LH14 seeds among fresh harvested (FS), after ripened (DS), and just-germinated stage (GS), the RNA-seqs of the six samples from three stages were performed by Illumina sequencing. In total, 44.5 to 63.4 million raw reads from the six libraries with an error rate of approximately 0.03% were generated, and the 44.1∽ 62.7 million of clean reads generated by removing low-quality sequences were selected for further analysis (Table 2). Among which, 81.11∽85% of reads were mapped to the reference genome (*Arachis duranensis*, https://www.peanutbase.org/home), and the reads uniquely mapped account for 78.34∽ 82.63%. The percentages of reads mapped into exon, intron, and intergenic region were 83.1∽87.5%, 1.3∽3.6%, and 11.1∽15.1%, respectively.

**Table 2.** Summary of the transcriptome sequencing data

| Sample_name | FS_1 | FS_2 | DS_1 | DS_2 | GS_1 | GS_2 |
| --- | --- | --- | --- | --- | --- | --- |
| Raw Reads | 59931664 | 44502832 | 56994704 | 63409368 | 50687486 | 51665824 |
| Clean reads | 59307780 | 44076312 | 56410796 | 62736078 | 50126196 | 50974284 |
| Q30 (%) | 93.56 | 93.72 | 93.70 | 93.46 | 93.18 | 91.92 |
| Total mapped | 48105991<br>(81.11%) | 36147297<br>(82.01%) | 45822510<br>(81.23%) | 50745769<br>(80.89%) | 42171822<br>(84.13%) | 43329793<br>(85%) |
| Uniquely mapped | 46684962<br>(78.72%) | 35267221<br>(80.01%) | 44419221<br>(78.74%) | 49145234<br>(78.34%) | 40939147<br>(81.67%) | 42120239<br>(82.63%) |
| Exon | 83.7% | 87.5% | 85.5% | 84.5% | 83.1% | 83.1% |
| Intron | 1.3% | 1.4% | 1.8% | 2.6% | 3.1% | 3.6% |
| Intergenic | 15.1% | 11.1% | 12.7% | 12.8% | 13.9% | 13.2% |

### Analysis of Differentially Expressed Genes (DEGs) at Different Stages

To investigate the major genes controlling dormancy release and germination of peanut seeds, the analyses of DEGs in the three sample pairs (DS vs FS, GS vs FS, and GS vs DS) were performed. A total of 3440, 2295, and 4657 DEGs were identified in above comparative pairs, respectively. There are 2169 up-regulated genes, and 1271 down-regulated genes between the after-ripened and the fresh-harvested seeds, and 1056 up-regulated and 1239 down-regulated ones between the just-germinated and the fresh-harvested seeds. Between the just-germinated and the after-ripened period, there are much more DEGs than those in other two comparisons, among them, the expression level of 2200 genes increased, and 2457 genes decreased. Of the total 5425 up-regulated and 4967 down-regulated DEGs, only 65 and 63 DEGs shared commonly across the three stages, and about 288, 728 and 65 enhanced DEGs, and 391, 629 and 63 reduced DEGs respectively overlapped in different combination of sample pairs, and 1881 and 880, 282 and 105, as well as 1472 and 1828 up-regulated and down-regulated genes were specifically found in DS vs FS, GS vs FS, and GS vs DS comparisons (Fig 2). It is interesting that between just-germinated seeds and fresh-harvest seeds, the morphologic changes were much obvious, while the number of DEGs was the fewest across the three pairwise.

**Fig 2.**
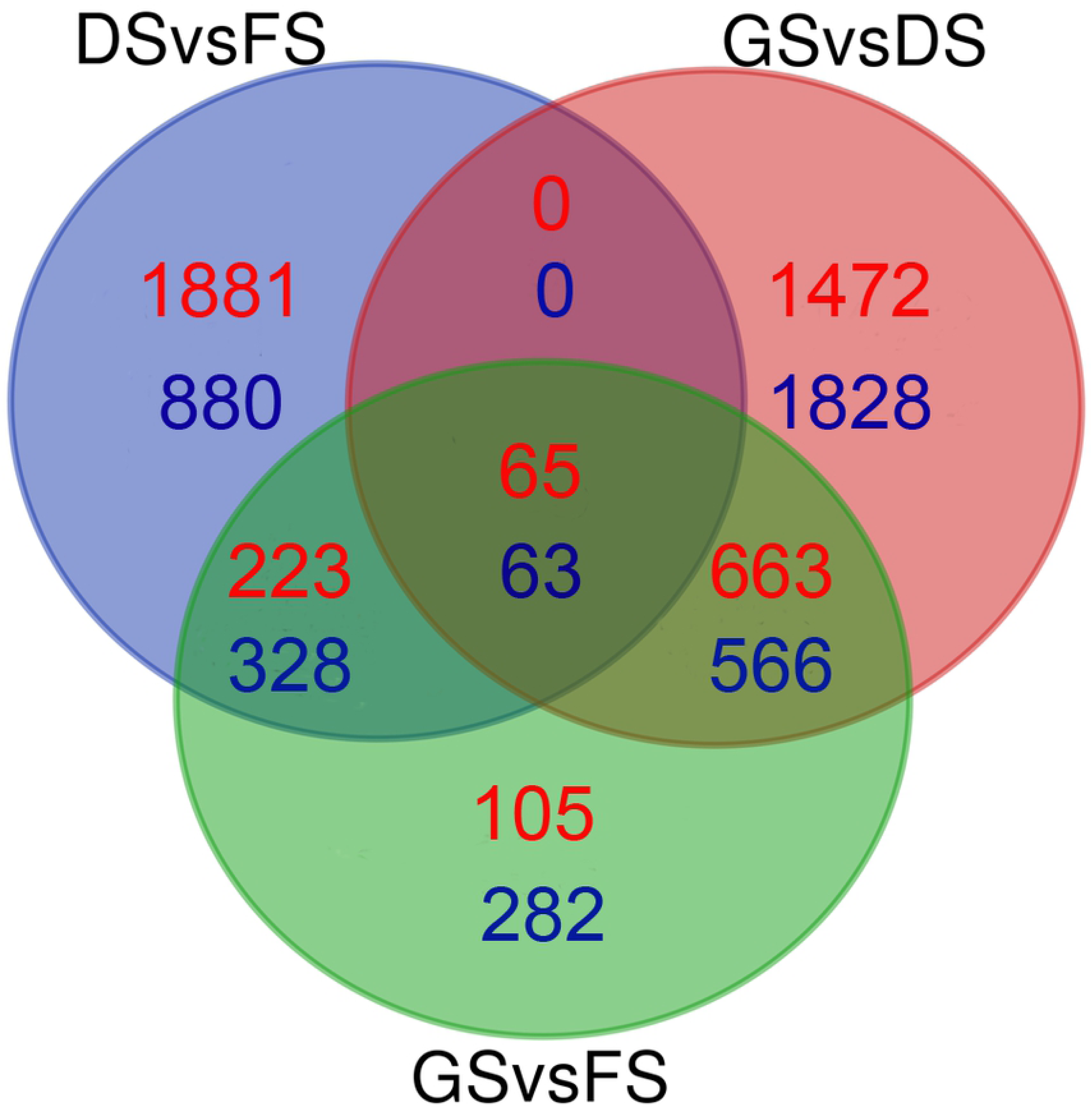
Venn diagrams of differentially expressed genes (DEGs) among fresh-harvest seeds (FS), dired seeds (DS), and just-germinated seeds (GS). Red counts represent up-regulated and blue counts are down-regulated between different comparisons.

### GO Functional Classification of DEGs at Different Stages

The Analysis of GO was performed to explore the biological processes related to dormancy release and germination of peanut kernels. In DS vs FS, the up-regulated DEGs were significantly enriched in 21 GO terms (corrected *p* < 0.05), in which the top seven terms were oxidation-reduction process (GO: 0055114, 241 genes), single-oganism biosynthetic process (GO: 0044711, 177 genes), organonitrogen compound metabolic process (GO: 1901564, 227 genes), cofactor metabolic process (GO: 0051186, 78 genes), small molecule metabolic process (GO: 0044281, 163 genes), pigment metabolic process (GO:0042440, 46 genes), and isoprenoid biosynthetic process (GO: 0008299, 23genes), while there were no down-regulated DEGs markedly enriched in GO terms. In this comparison pair, the majorities of the improved DEGs focused on the molecular function of oxidoreductase activity (Fig 3A). Compared GS to FS, only the down-regulated DEGs were grouped into one GO term, that is the embryo development process (GO: 0009790), in which 7 genes out of 20 genes located in the reference genome was detected (Fig 3B). In GS vs DS, the up-regulated DEGs were remarkably enriched in 19 GO terms, which involve in several biological regulation and cellular process including regulation of gene expression (GO:0010468, 184 genes), regulation of RNA biosynthetic process (GO:2001141, 177 genes), protein phosphorylation (GO:0006468, 138 genes), cellular protein modification process (GO:0006464, 183 genes), and so on, and among them, the crucial parts mainly execute the following molecular functions including nucleic acid binding transcription factor activity, ubiquitin-protein transferase activity, and so on (Fig 3A). In this comparative pairwise, the down-regulated DEGs mainly grouped in 11 GO terms, among which the majorities were detected in the upregulated DEGs of DS vs FS, including oxidiation-reduction process (GO: 0055114, 291 genes), single-organism biosynthetic process (GO: 0044711, 201genes), pigment metabolic process (GO: 0042440, 51genes), cofactor metabolic process (GO: 0051186, 83 genes), and etc., and other enrichment terms related to metabolic process, mainly including carbohydrate derivative metabolic process (GO:1901135, 125 genes) and carbohydrate metabolic process (GO: 0005975, 161 genes) (Fig 3B).

**Fig 3.**
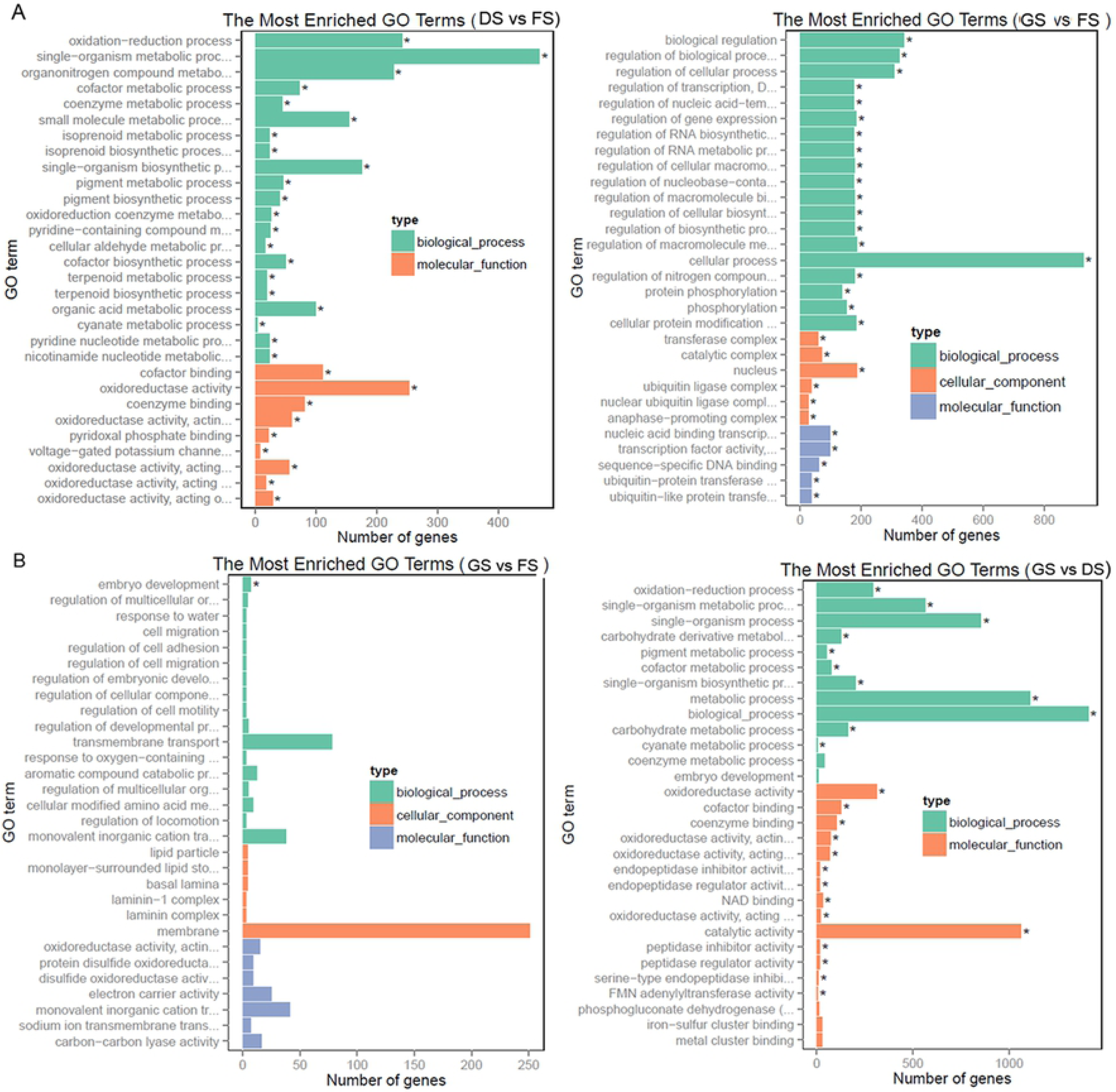
The histogram of DEGs functional classification between DS and FS, GS and DS, and GS and FS using GO annotation. A. The classification of up-regulated DEGs; B. The classification of down-regulated DEGs

### Rapidly Reactivation of Energy Metabolism during After-ripening

A large number of enzymes and mRNAs involved in the major metabolic pathways including energy production pathways store in maturated dry seeds, preparing for germination of seeds and seedling establishment [34–37]. During a period of dry storage, which seeds undergo the period of after-ripening, seed dormancy is gradually alleviated [35]. In our study, the KEGG pathwayanalysis showed that during the after ripening of peanut seeds, many genes involved in oxidative phosphorylation (S1 Table, gmx00190, 48/243genes, corrected *p*=2.48e-05), glutathione metabolism (gmx00480, 28/155 genes, corrected *p*=0.015), and carbon metabolism (S2 Table, gmx 01200, 59/447 genes, corrected *p*=0.021), were significantly increased.

Mitochondria is the important place where respiration takes place, during which electrons are transferred from electron donors to acceptors to produce reactive oxygen species such as superoxide O_2_- and hydrogen peroxide H_2_O_2_, and the energy in the way of adenosine triphosphate (ATP) are released [38]. In our study, during the after-ripening of peanut seeds, the enzymes encoded by the up-regulated crucial DEGs significantly associated with the complexes I to V of the electron transport chain in oxidative phosphorylation, including NADH dehydrogenase (ND) subunit 1, 2, 4, 4L, 5 and 6, and NADH dehydrogenase (ubiquinone) iron-sulfur (Ndufs) subunit 1, 2, 7, 8 and flavoprotein 2 (Ndufv2)from complex I (NADH-coenzyme Q oxidoreductase), succinate dehydrogenase (ubiquinone) iron-sulfur subunit 2 (SDHB2) from complex II (succinate-Q oxidoreductase), ubiquinol-cytochrome c reductase iron-sulfur subunit (ISP), and cytochrome b subunit (Cytb), cytochrome c1 subunit (Cyt1) and ubiquinol-cytochrome c reductase subunit 7 (QCR7) from complex III (cytochrome bc1 complex), cytochrome c oxidase (COX) subunit 1, 2 and 3 from complex IV, and the different kind of ATPase from complex V(F-type H+-transporting ATPase subunitα, subunit a, b and g, V-type H+-transporting ATPase subunit B, D, E, G, H and 21kDa proteolipid subunit) (Figs 4A and 4B, S1 Table). The qRT-PCR results also verified that the expression of some key genes involved in complexes I to V of the electron transport chain increased at dry seed stage (Fig 5A).

**Fig 4.**
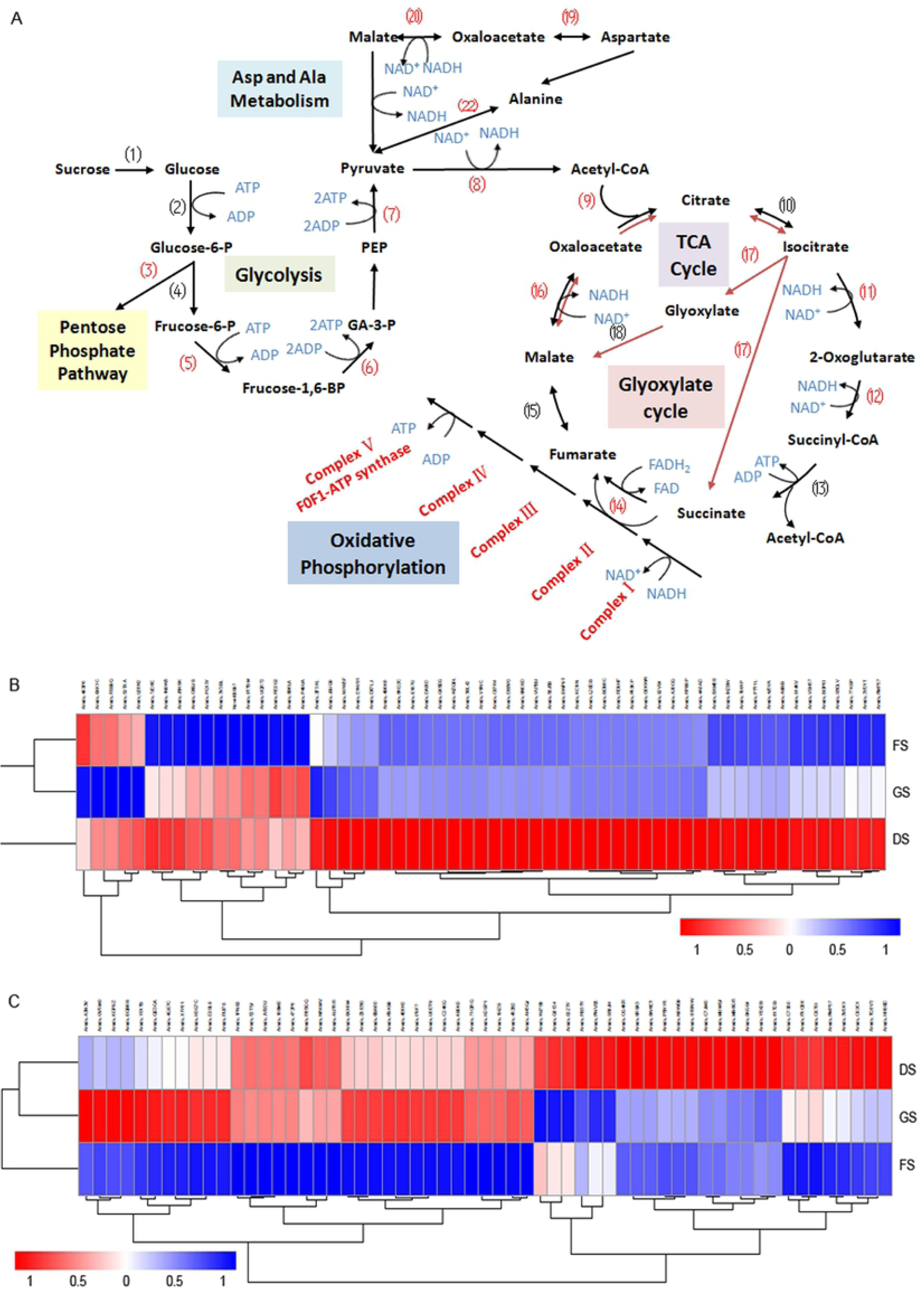
Differences of gene expression between DS and FS stage in several metabolic pathway. A. The majority of up-regulated genes represented in the KEGG pathway involved in glycolysis, tricarbooxylic acid cycle (TCA), glyoxylate cycle, ASP and ALA metabolism, and oxidative phosphorylation. (1) Invertase; (2) Hexokinase (HK); (3) Glucose-6-phosphate dehydrogenase (G6PDH); (4) Pyrophosphate-dependent phosphofructokinase (PFP) / Diphosphate-fructose-6-phosphate1-phosphotransferase;(5) Phosphofructokinase (PFK); (6) Fructose-biphosphate aldolase (FBA); (7) Pyruvate kinase(PK); (8) Pyruvate dehydrogenase complex; (9) Citrate synthase (CSY); (10) Citrate hydro-lyase and Citrate hydrooxymutase; (11)Isocitrate dehydrogenase (IDH); (12) 2-oxoglutarate dehydrogenase (OGDH); (13) Succinyl-CoA:acetate CoA transferase/SSA-CoA synthetase; (14) Succinate dehydrogenase (SDH); (15) Fumarate hydratase; (16) Malate dehydrogenase (MDHm); (17) Isocitrate lyase (ICL); (18) Malate synthase (MSY); (19) Aspartate aminotransferase (AspAT); (20) Malate dehydrogenase (MDHc); (21) NAD-dependent malic enzyme 2 (NAD-ME2); (22) Alanine aminotransferase (AlaAT). The numbers in parentheses marked in red color represent the up-regulated genes encoding the key enzymes in related pathway. B. Heatmaps of the 59 DEGs among FS, DS and GS stage in oxidative phosphorylation pathway; C. Heatmaps of the 59 DEGs among FS, DS and GS stage in carbon metabolic pathway

**Fig 5.**
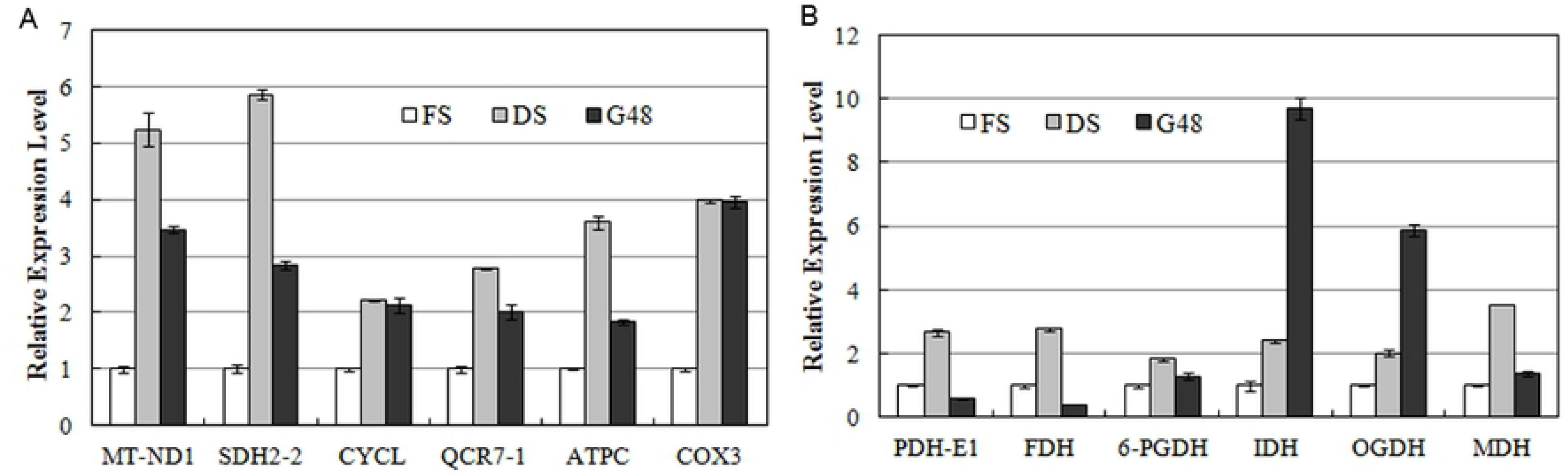
Analysis of mRNA transcription level of several DEGs between DS and FS stage by real-time fluorescent quantitative RT-PCR. A. The DEGs related to the electron transferring chain in oxidative phosphorylation pathway; B. The DEGs involved in dehydrogenation reaction in glycolysis and TCA

The reserves such as starch, lipid, and protein stored in plant seeds, act as the source of carbon and nitrogen, which are degraded and mobilized during filial germination and seedling establishment [39]. Except for these reserves, lots of metabolites including sugars, organic acids, polyols, amino acids, and some fatty acid-related compounds are accumulated in maturation seeds, which also as the energy storage, could provide certain metabolites rapidly to recover the corresponding metabolic pathways before mobilization of storage reserves [6]. Our results of KEGG pathway analysis showed that mobilization of some metabolites is rapidly promoted during the after-ripening of peanut seeds with the shorter period of dormancy. Lots of genes associated with glycolysis, tricarboxylic acid (TCA) cycle (also named as citrate cycle), and glyoxylate cycle were significantly up-regulated during this stage (Fig 4, S2 Table). Among them, twenty-one genes encoding different dehydrogenase, including pyruvate dehydrogenase complex, isociteate dehydrogenase (IDH), 2-oxoglutarate dehydrogenase, succinate dehydrogenase, and malate dehydrogenase (MDH), and etc., account for 1/3 of up-regulated genes, which by a series of oxidation reaction of intermediates in glycolysis pathway and TCA cycle, catalyze one pyruvate molecule to produce CO_2_, one molecule of ATP, and four NADH and one FADH_2_ molecules [40]. Six genes out of them had been confirmed to be up-regulated in dry seeds by qRT-PCR (Fig 5B). TCA cycle also could oxidize the intermediates of amino acids by a transamination reaction [41]. In cotyledons of soybean and pea, aspartate aminotransferase participate in this catabolic reaction to generate an intermediate of TCA [26]. The transcriptional level of four genes encoding aminotransferase (Aspartate aminotransferase and Alanine aminotransferase 2 from mitochondria, Serine-glyoxylate aminotransferase, and Phosphoserine aminotransferase 1 from chloroplast) was remarkably elevated in our results (Figs 4A and 4C, S2 Table). The resulting NADHs and FADH_2_ molecules enter into electron transport chain and are further oxidated to produce energy by oxidative phosphorylation. Total 12 ATP molecules are generated by TCA cycle [41]. In addition, expression of several genes encoding glyceraldehyde-3-phosphate dehydrogenase also improved, which catalyze the oxidation and phosphorylation of glyceraldehyde-3-phosphate to produce 1,3-bisphospho-D-glycerate in glycolysis.

Therefore, after a period of storage, the intermediates stored in peanut dry seeds are rapidly mobilized by glycolysis, TCA cycle, and glyoxylate cycle, and some transamination process; and the electron transport chain accompanying with respiration has been reactivated to provide ATP for mobilization of other reserves and seed germination. During this period, ROSs as by-product also accumulated in seeds, which are considered to associate with the status transformation from dormancy to non-dormancy [4, 5].

### Multiple Pathways of Plant Hormone Signal Transduction during Seed Germination

During germination of peanut seeds, some down-regulated genes were classified into oxidative phosphorlation pathway (41 genes, corrected *p*=0.0025), and dozens of the up-regulated genes were related to plant hormone biosynthesis and signal transduction (Figs 4B and 6A, S1 and S3Tables). Indole-3-acetic acid biosynthesis is the necessary trigger for seed germination [6]. The majority of components in auxin signal pathway, including auxin transporter-like protein AUX1 which is an auxin influx carrier, the F-box protein TRANSPORT INHIBITOR RESPONSE 1 (TIR1), AUXIN RESPONSE FACTOR (ARF), probable indole-3-acetic acid-amido synthetase GH3, SAUR family protein, and AUX/IAA family proteins, were significantly improved in this period. By qRT-PCR, the expression level of several crucial genes related to IAA signaling also had been confirmed to be induced in germinated seeds (Fig 7A). Lots of genes involved in brassinosteroid biosynthesis and signal transduction were also markedly increased, such as cytochrome P450 90B1 and 90A1, steroid 5-alpha-reductase DET2, brassinosteroid receptor BRI1 (brassinosteroid insensitive 1), BRI1-associated receptor kinase BAK1, and brassinosteroid resistant BZR1 and BRZ2; while one *BIN2* gene was down-regulated significantly. In addition, the up-regulated genes also included some GA and ABA signal transduction genes, for example, gibberellin receptor GID1 (GA-insensitive dwarf 1), F-box protein GID2, abscisic acid receptor PYR/PYL, protein phosphatase 2C (PP2C), and serine/threonine-protein kinase SnRK2 (S3 Table).

**Fig 6.**
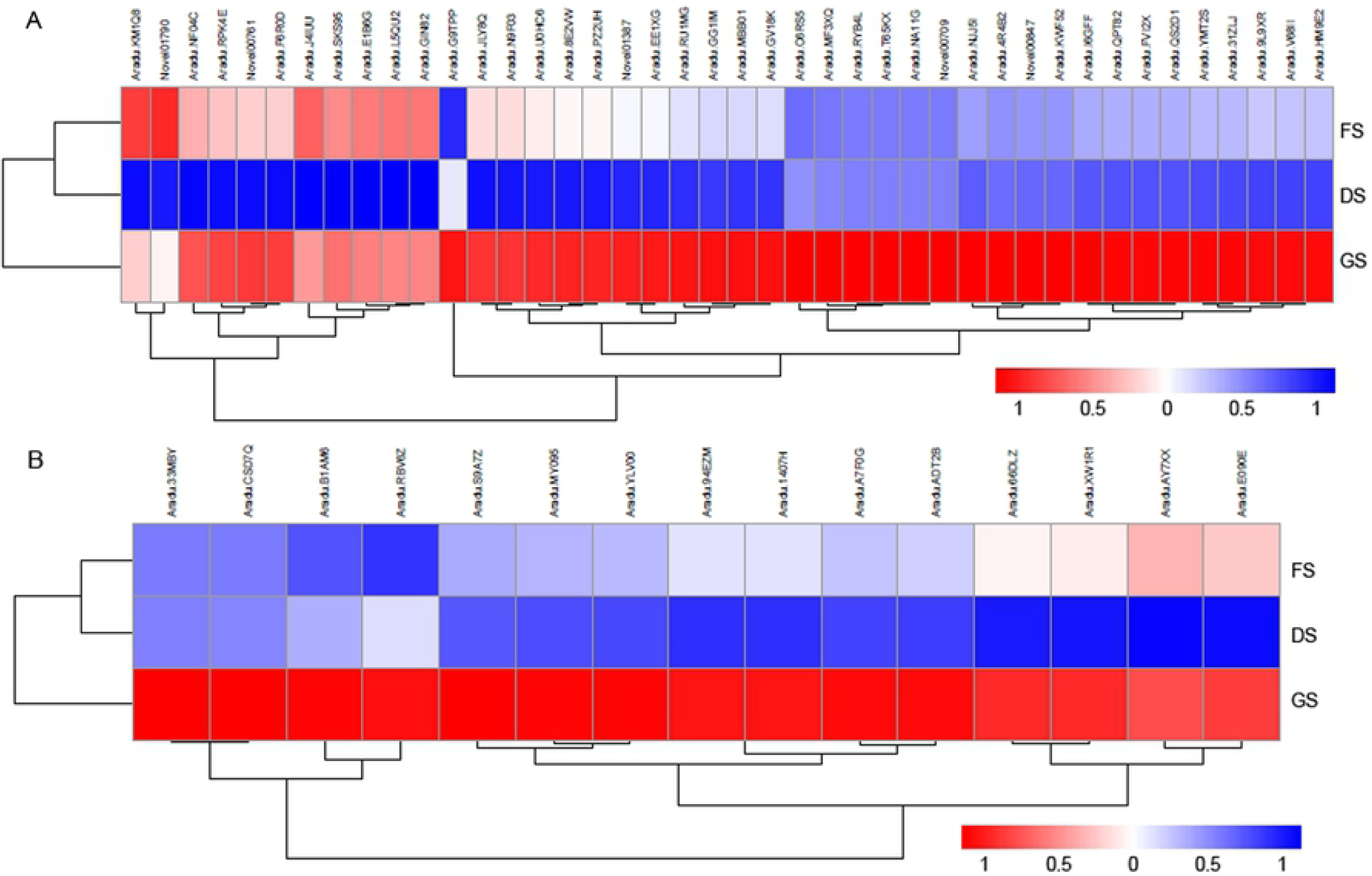
The signaling pathway of phytohormone including GA, BR, auxin, and so on is enhanced in the germinated seeds. A. Heatmaps of the 42 DEGs among FS, DS and GS stage in phytohormone signaling pathway B. Heatmaps of the 15 DEGs among FS, DS and GS stage in protein ubiquitin degradation pathway

**Fig 7.**
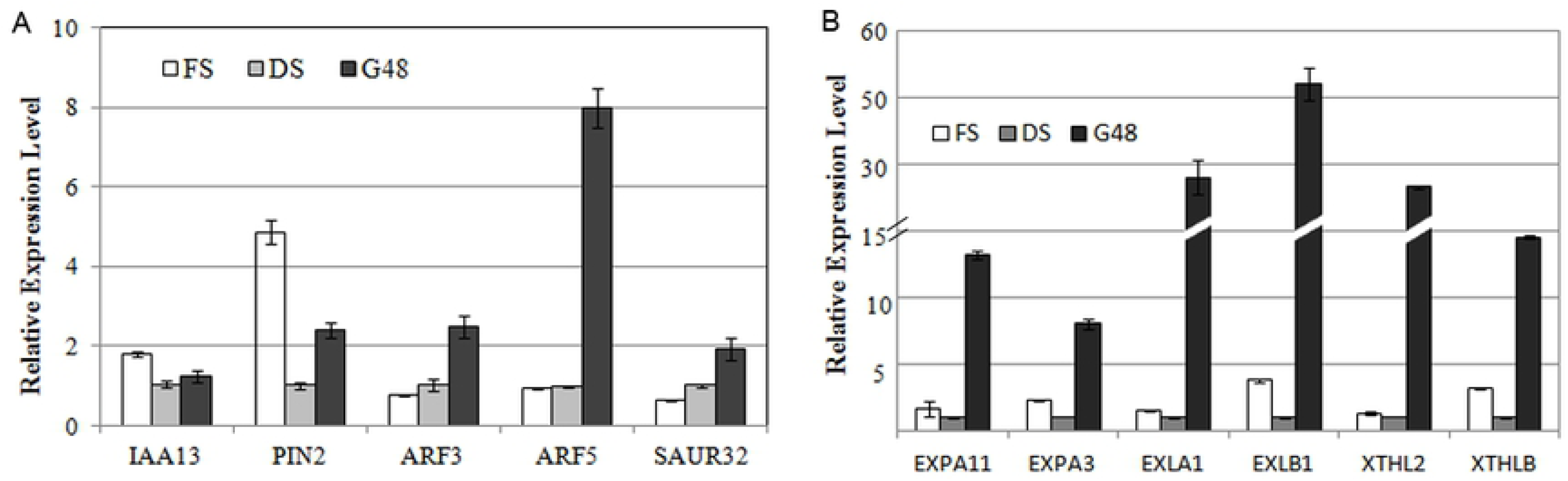
Analysis of mRNA transcription level of several DEGs between GS and DS stage by real-time fluorescent quantitative RT-PCR. A. The DEGs involved in auxin signal transdution pathway; B. The DEGs related with cell elongation and cell wall remodoling

Ubiquitin-mediated proteolysis plays an important regulatory role in hormone signaling [42], during which the proteins are polyubiqintinated by a reaction cascade that consist of three enzymes, named E1 (ubiquitin activating enzyme), E2 (ubiquitin conjugating enzyme), and E3 (ubiquitin ligase), and are degraded by 26s proteasome. Our research found that some genes encoding E1, E2, and E3 were significantly improved during seed germination, which include ubiquitin-like 1-activating enzyme E1 A (UBLE1A), ubiquitin conjugating enzyme (UBE2C, UBE2D, UBE2I, and UBE2O), and Cullin (Cul1), adaptor protein Skp1 (S-phase kinase-associated protein 1) belonging to SCF complex of multi subunit RING-finger type E3, and so on (S4 Table, Fig 6B).

### The Distribution and Content of Hormones in Different Stages of Seed

Phytohormone plays an important role in determining the physiological state of the seed and in regulating the germination process [11,37]. Therefore, in order to understand the crucial roles of different hormones in seed dormancy, dormancy release and germination. The contents of ABA, GA, BR and IAA in different developmental stages of seed were detected, and the whole seed was divided into three parts (CO, HR and PL) for exploring the differential distribution of hormones.

ABA is a positive regulator of dormancy induction and a negative regulator of germination, while GA counteracted with ABA release dormancy and promoter germination. Our detection results found that ABA contents in every part of fresh-harvest seeds displayed higher level, and ABA level decreased in the HR part of dried seeds, while significantly increased in the CO and PL part. ABA contents in CO and PL rapidly declined during the early phase of seed imbibitions, and kept in a constant level after imbibition for 28h, while ABA content in HR slightly rose during the early stage, and began to decrease after 28h. In fact, seeds of imbibition for 28h and 52h have lower ABA level (47.48 and 52.74 ng/g.FW). GA_3_ levels in HR of all development stages, from 10.53 ng/g.FW to 14.94 ng/g.FW, were much higher than those in CO and PL, that was lowest in dried seeds, and was highest in imbibition seeds for 4h and dropt with the duration of germination time. In CO, GA_3_ content decreased accompanying water loss to dried seeds, and also maintained a lower level in imbibition for 4h, and then increased with the prolongation of imbibition time and reached the top level in imbibition for 28h, and then slightly reduced. In PL, there was the highest GA_3_ content in dried seeds, and during the procedure of germination, GA_3_ contents kept the lower level (Fig 8A). Previous researches indicated that it is likely that the ABA: GA ratio, and not the absolute hormone amounts, regulates dormancy release and germination. Dormancy maintenance depends on higher ABA: GA ratio, while dormancy release involves a net shift to increased GA biosynthesis and ABA degradation resulting in lower ABA: GA ratio [7, 11]. Therefore, the ABA: GA ratios in three seed parts of all development periods were assessed. The results showed that low ABA: GA ratios were maintained at all stages in HR, while in PL, high ABA: GA ratios except for at dried seeds; and in CO, ABA: GA ratio in dried seeds was much higher than those in other periods, and with the duration of germination, ABA: GA ratios sharply dropped from 71.08 to 3.82 (Fig 8A).

**Fig 8.**
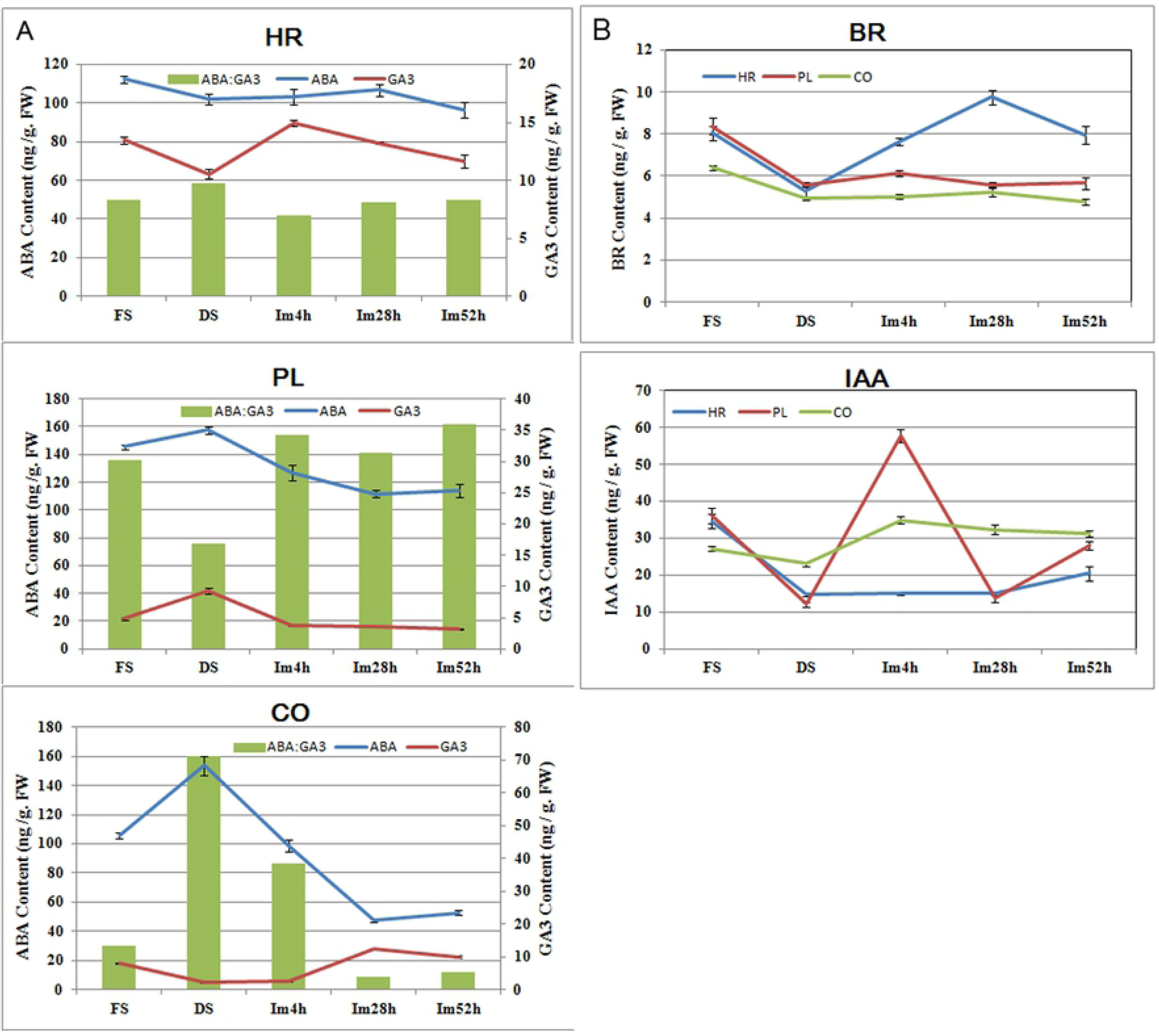
The phytohormone contents of different section from peanut seeds at different developmental stages. A. ABA and GA content, and the ratio of ABA to GA in fresh-harvested seeds, dried seeds, and the seeds during imbibition for 4, 28 and 52hr; B. BR and IAA content in fresh-harvested seeds, dried seeds, and the seeds during imbibition for 4, 28 and 52hr. HR: hypocotyl and radicel; PL: Plumule; CO: Cotyledon

BR and GA as the positive regulators, counteract the inhibitory action of ABA, promote cell elongation and seed germination in parallel way [11]. Our results found that in CO and PL, BR contents slightly changed during seed germination, while in HR, BR level was lowest in dried seed, and then enhanced significantly when seed began to suck up water, and reached the peak in imbibitions for 28h following a distinct decrease (Fig 8B).

By now, very little is known about the role of auxin during seed germination. However, some studies found that auxin maybe interact with GA and resulting in the change of IAA synthesis and transport during germination [11,43]. Our detection results found that in any part of dried seeds, the IAA contents were much lower; and then during imbibiting for 4h to 28h, IAA kept the lowest level in HR, while sharply rose up and went down in PL; and the accumulation of IAA enhanced in HR and PL with the duration of imbibition time. In CO, IAA level significantly increased at early stage of water uptake, and then dropped down little by little till imbibition for 52h (Fig 8B).

## Discussion

### Dormancy Alleviation of LH14 Dry Seeds Associated with the Rapid Resumption of Multiple Metabolic Processes during After-ripening

Seed dormancy is established during seed maturation, and dormancy loss of maturated seeds can take place through a period of dry storage (so-called as AR), through moist chilling, or through seed coat scarification. Different species or same species living in variant natural habitats evolve to have different dormancy adaption for environmental conditions. For instance, Landsberg *erecta* (Ler) or Columbia (Col) ecotype of Arabidopsis have a low level of dormancy, while the seeds of Cape Verde Islands (Cvi) ecotype show a strong dormancy [10]. AR is a specific developmental procedure, which broaden or increase sensitivity of perception of seeds to environmental conditions for promoting germination, and simultaneously decrease or narrow sensitivity of perception in repressing germination [7,44]. The duration time of AR procedure is much different among different species, some go through several days, and others last for several months or more [10,15,25]. Although after-ripened seeds have been provided with germination ability, their germination must meet appropriate environment including suitable temperature, humidity, and etc., or else seeds will reenter the second dormancy. Physiological status of dry seeds seems to be quiescent, however, abundant changes in gene expression have been found in dry seeds compared to those in dormant seeds, which trigger by AR [45–47]. It was shown that some genes associated with storage mobilization, cell wall modification were highly expressed in after-ripened seeds to dormant seeds of Arabidopsis and wheat [45,46]. Although transcriptional profiles and molecular mechanism underlying dormancy release by AR are conserved between species, there are their own unique regulatory mechanisms in different species [46,48]. In this study, seeds of LH14 display the obvious AR period during dry storage despite easily losing their dormancy. The results of transcriptomic analysis of dormant and AR peanut seeds indicated that lots of genes (3440 genes) were differentially expressed, of which the majority involved in multiple metabolic processes including the oxidative phosphorylation, carbohydrate metabolism, and glutathione metabolism modules. After-ripened seeds with dormancy have made the necessary preparation for germination and seedling establishment, in which the mobilization of reserves and energy production plays the crucial roles. Acetyl- CoA, producing through a transacylation reaction in glycolysis, is central for energy metabolism, which is oxidized via TCA cycle in mitochondria, and in glyoxysome, which also is an intermediate in the conversion of fatty acids to carbohydrates [26,49]. TCA cycle also could oxidize the intermediates of amino acids by a transamination reaction [26,41]. In peanut dry seeds, we found that large amounts of key genes associated with glycolysis, TCA and glyoxylate cycles, and amino acid metabolism, were highly transcribed. It was suggested that the stored soluble carbohydrates, fatty acids, amino acids and other intermediates could be rapidly utilized by resuming of several metabolic pathways, and an early activation of oxidative phosphorylation by electron transport chain could produce large amounts of ATP to supply the following procedure. Some studies in soybean, Arabodopsis and sugarbeet indicated that glycolysis, fermentation, TCA glyoxylate cycles and the oxidative pentose phosphate pathway (OPPP) are quickly activated by AR upon imbibitions to supply energy for germination [15,26,34,37]. Therefore, in fact like peanut dry seeds, several catalytic procedures involved in energy metabolism might be rapidly resumed during AR of dry seeds in other dicot species; merely peanut dry seeds with non-deep dormancy are more sensitive to AR. The key genes corresponding to these pathways maybe still express in a higher level during the early phase of peanut seed germination, while majority of them display the down-regulated expression patterns during the later period of germination and testa breaking.

### Coordination of Hormone Signal Transduction Nets Plays a Key Role in Radicle Protrusion

During the period of radicle protrusion, breakage of seed testa and hypocotyl elongation is the major visible characters. Various phytohormones and environment signals take part in the regulation of this procedure.

The anagonistic effects of GA and ABA on seed dormancy breakage and germination have been clarified in many monocot and dicot species [1,7,10,11,50,51]. During the procedures of seed maturation, inception and maintenance of dormancy, dormancy release, and seed germination, the contents of ABA and GA in seeds, and the sensitivity of seeds to them, have much complex dynamic relationships. Although GA accumulation correlates with dormancy release and germination, GA treatment alone apparently does not satisfy the conditions of seed germination. A reduction in ABA levels is prerequisite before GA contents and sensitivity begin to increase [50]. In fact, maintaining of dormancy requires a higher ABA: GA ratio, while dormancy breakage and germination depends on the conversion of increasing GA biosynthesis and ABA degradation resulting in lower ABA: GA ratio [7,11,50]. In the present study, we found that the amount of ABA in cotyledon sharply dropped down at the onset of dry seeds sucking water, and maintained relative constant level during the late phase of imbibition; while GA level kept to increase after sowing in water for 4h, and hold on steady level after imbibition for 28h. Clearly, prior to the enhancing of GA content, the ABA content has dropped to a lower level, and the resulting lower ABA: GA ratio in peanut cotyledon is beneficial for the radicle breaking through testa. We didn’t found any genes involved in GA synthesis remarkably improved in germinated seeds, but found that four significant up-regulated genes participated in GA signal transduction, two of them encode GA receptor GID1 and others encode F-box protein GID2 which is the major component of the SCF^SLY1/GID2^ ubiquitin E3 ligase complex. However, during this period, the expression of a *DELLA* gene (gmx:547719), repressing GA signal transduction cascade, was significantly down-regulated. In GA signaling module, GA, receptor GID1 together with repressor DELLA form a GA-GID1-DELLA complex. When bioactive GA level raises, GID1 combining with GA occurs the conformational change, and DELLA is recruited to the SCF^SLY1/GID2^ ubiquitin E3 ligase complex for poly-ubiquitination and subsequent degradation by the 26S proteasome, relieving the suppressive effects on downstream GA-responsive genes [20,52]. We also found some genes involved in ubiquitin mediated proteolysis pathway are markedly up-regulated in germinated seeds; out of them included one gene encoding the SCF ubiquitin ligase complex protein. It is implied that the sensitivity of GA perception maybe increase on the initial of radicle emergence. Furthermore, GA triggers seed germination by removing the mechanical restraint of the seed coat and endosperm, during which the expression of some expansin (EXP) and and *xyloglucan endotransglycosylases* (XETs/XTHs) family members are induced [37,50,53–56]. Our results indicated that the expression levels of eight peanut *EXP* genes [including five *α-expansin* (*EXPA*), one *expansin-like A* (*EXPLA*), and two *expansin-like B* (*EXPLB*)] and six *XTH* genes significantly up-regulated at the stage of seed germination compared to those in dry seeds (Fig. 7B).

Both GA and BR stimulate seed germination by different regulatory mechanism [50,57,58]. Although both GA and BR can induce the expression of cell elongation- or cell wall organization-related genes including *EXP*s, but these hormones promote the expression of distinct family members. BR is considered to promote seed germination by directly improving the growth potential of embryo in a GA-independent manner [50,58]. When BR content is high, BR is perceived and bound by BRI1, and activated BRI1/BAK1 kinase complex, and thereby the downstream repressors of BR signaling, including the GSK3-like kinase BIN2, are inhibited; the inhibition of BIN2 results in the accumulation of unphosphorylated BZR1/2 family transcription factors that regulate BR-target gene expression [12,59–62]. In the present study, the expression levels of some transcripts associated with positively regulating BR biosynthesis and signaling pathway were remarkably up-regulated during germination of peanut seeds, those include *DET2*, *BRI1*, *BAK1*, *BRZ1*, and so on; while one kinase *BIN2* gene (gmx:100802451), negatively regulating BR signal transduction, was markedly down-regulated. In the meanwhile, we found that the significant increase of BR levels only took place in the hypocotyls and radicles of imbibed peanut seeds. It is suggested that the elevation of BR content in the HR section of peanut imbibed embryo was rapidly perceived by BRI1, and the BR signal transduction cascade was subsequently initiated, resulting some genes required for cell elongation were activated in the imbibed hypocotyls and radicles until seed germination finished. This is consistent with the conclusion that increase of BR signaling intensity improves the status of seed germination, and increases the length of hypocotyls [60,63,64].

Auxin is a major hormone associated with plant morphogenesis, which is also essential for promoting hypocotyl elongation and seed germination [64–67]. The SCF^TIR1^ –auxin – AUX/IAA complex is the central component of Auxin signaling model, in which auxin triggers ubiquitination and degradation of the AUX/IAA family proteins to derepress the inhibition of ARF transcription factors, and subsequently to promote the expression of some auxin-responsive genes [68]. In this study, at the radicle protrusion time point, the expressions of a large number of genes involving auxin signal transduction (AUX1, TIR1, CULLIN, AUX/IAAs, ARFs, GH3, SAUR) were substantially increased; the transcription levels of two *PIN2* genes (encoding auxin efflux carrier) were also significantly produced. While at this time, no obvious expression changes were observed in genes specifying for auxin biosynthesis. Taken together, it was implied that during the later stage of peanut seeds germination, auxin transport is active and the auxin signaling pathway plays the important regulatory roles. Auxin distribution in Arabidopsis young seedlings is imbalance, a higher level in root apex and cotyledon; and its polar transport associates with root morphogenesis and gravitropism, inhibiting auxin transport results in aberration of root gravitropism and elongation [69,70]. We found that during the earlier stage of imbibitions, IAA content in CO and PL section of peanut seeds rapidly increased up to a higher level, especially in PL section, while in HR section it kept constant level until approaching germination, and then started to rise slowly, indicating that the increase of IAA level maybe relate to the elongation of embryo axis (HR section) during germination.

BR- and auxin-mediated cell elongation is interdependent, and this synergism doesn’t depend on the level of hormone biosynthesis [67]. The crosstalk of auxin and BR signals is found to converge on the regulation of ARF transcription factors, which is downstream from BZR1 and AUX/IAA proteins and trigger the expression of some auxin-response genes with ARFAT motif (TATCTC) in the promoters [65,67]. Walcher and Nemhauser (2012) found that BZR2 and ARF5 could bind to the 5′flanking region of *SAUR15* gene that is activated by both auxin and BR [71]. Oh et al., recently clarified a central regulation cassette in regulating hypocotyl cell elongation that auxin, BR, GA, light, and temperature signals were integrated together. In this module, BZR1and light –responsive factor PIF4 co-regulate hypocotyl cell elongation by interacting with specific ARFs such as ARF6, ARF8 and etc. However, GA signaling pathway in regulating cell elongation is converged through removing the DELLA proteins inhibition to specific ARFs, together with BZR1 and PIF4, to promote the expression of some ARF target genes [64]. At the germination time-point of peanut seeds, the homologous genes of Arabidopsis *ARF3*, *ARF5*, *ARF8*, *ARF18* and *ARF32* were significantly expressed up-regulatedly. However, among them, which is the central one or ones in integrating auxin signal with BR and GA signals? It needs to verify by experiments. Several studies indicated that seed germination procedure involve in the regulation of cell expansion and cell wall organization, during which the expression levels of some *XTH*s and *EXP*s genes improve remarkably[37,53–56,65,71]. Recent studies found the 5′flanking region of some *XTH*s, *EXP*s and other auxin-responsive genes including *IAAs*, *SAURs* and *GH3s* contain the TGTCTC or its inverse element GAGACA that is the binding site of some ARFs [64,65,67,71]. Our results also found that lots of genes mentioned above expressed at a high level in just-germinated seeds, suggesting that some specific ARFs together with other transcription factors regulating cooperatively by GA, BR, auxin, and other signals modulate the ARF target gene expression and promote breakage of peanut seed testa and hypocotyl elongation (Fig. 9).

**Fig 9.**
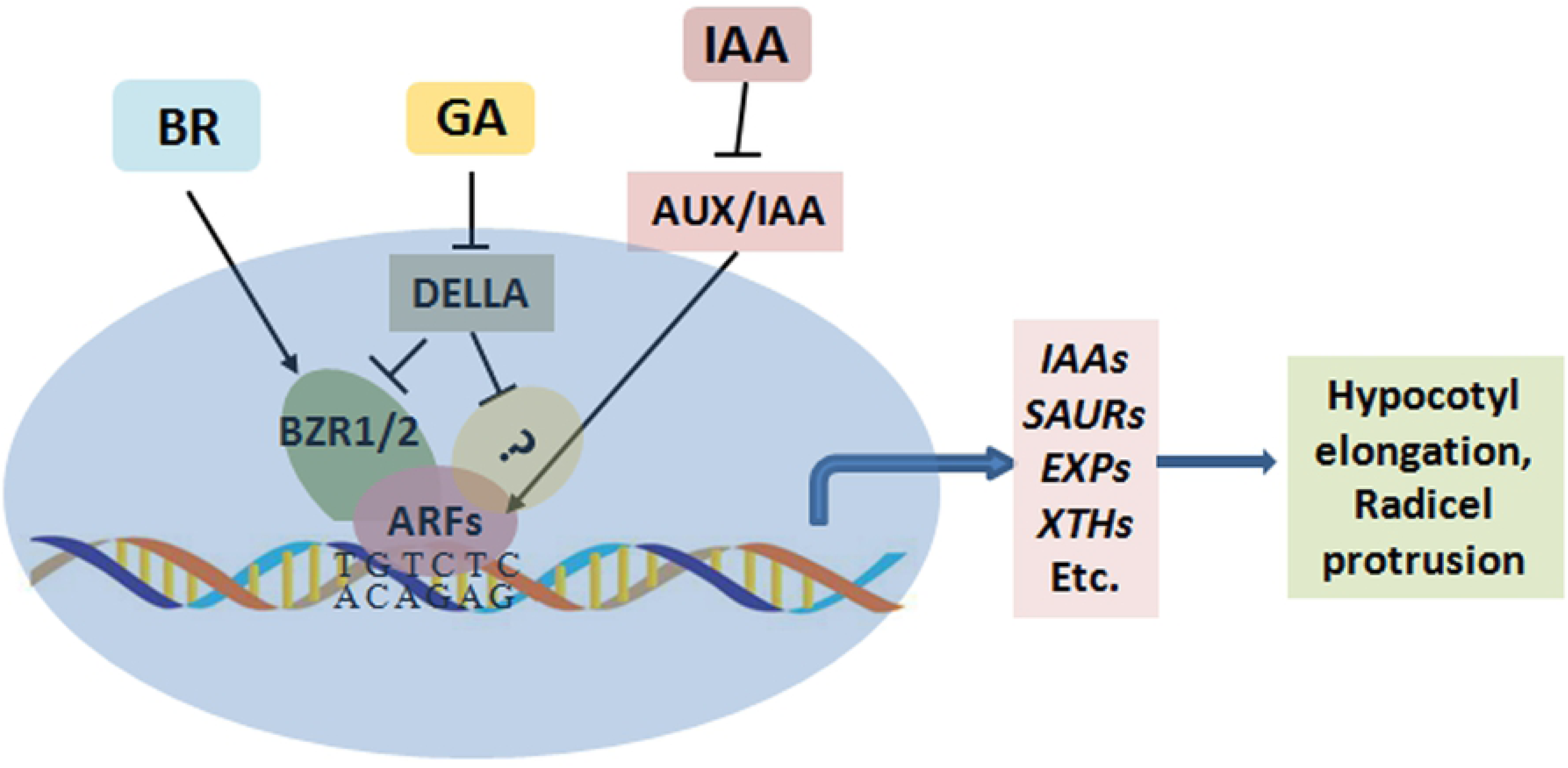
Model sketch of coordinating effects of hormone GA, BR and auxin signaling on hypocotyl elongation and radicel protrusion

## Supporting Information

**S1 Table DEGs in OP pathway**

**S2 Table DEGs in CM pathway**

**S3 Table DEGs in phytohormone signaling pathway**

**S4 Table DEGs in UB pathway**

## Author Contributions

LS, ZL, SW designed the experiments and contributed to the writing of the manuscript. PX, GT, WC, JZ performed the qRT-PCR verification and the detection of phtohormone. GC, CM performed bioinformatic analysis. LS, PL drew the diagram of metabolic pathways. All authors revised the draft and approved the final manuscript.

